# Prenatal alcohol exposure dysregulates the expression of clock genes and alters rhythmic behaviour in mice

**DOI:** 10.1101/2025.05.28.656556

**Authors:** Maria Reina-Campos, Ines Gallego-Landin, Mireia Medrano, Olga Valverde

## Abstract

Foetal Alcohol Spectrum Disorders (FASD) refer to a range of adverse physical, behavioural, and cognitive effects caused by perinatal alcohol exposure. While cognitive impairments are well-documented, FASD has also been associated with sleep disturbances and circadian rhythm disruptions. This study aimed to examine the effects of perinatal alcohol exposure on circadian rhythms at behavioural and gene expression levels across two developmental stages (adolescence and adulthood) in both, male and female mice. Using a validated prenatal and lactation alcohol exposure (PLAE) protocol, we assessed circadian patterns of locomotor activity under free-running conditions and spatial memory performance during adolescence and adulthood. Additionally, we evaluated the impact of PLAE on circadian expression of clock and non-circadian genes involved in neurotransmission across key brain regions, including the medial prefrontal cortex and hippocampus. PLAE altered circadian rhythmicity and impaired spatial memory. Gene expression analyses revealed disrupted oscillatory patterns in clock genes and in genes related to plasticity and cognition, including those from the expanded endocannabinoid system (e.g. *Cnr1*, *Dagla*, *Faah*) and other neurotransmitter systems (e.g. *Oprm1*, *Slc17a8*, *Drd1*, *Gabra1*). These findings underscore the impact of early alcohol exposure on biological rhythms and neurobehavioral function, highlighting circadian dysregulation as a contributing factor to FASD.

## INTRODUCTION

Foetal Alcohol Spectrum Disorders (FASD) refers to the range of adverse physical, behavioural, and cognitive effects caused by perinatal alcohol exposure (1). This condition can manifest as a range of symptoms which significantly differ among individuals. Among them, cognitive impairments are the most prevalent and disabling including intellectual capacity, executive functioning, motor performance, attention, learning, and memory (2). Currently, the global prevalence of FASD is estimated to be 0.77% of children and youth worldwide (3). However, it is widely accepted that prevalent rates are underestimates since its diagnosis relies on the history of maternal alcohol consumption during pregnancy and the assessment of the patient’s neurobehavioural profile. The lack of clear biological markers complicates detection, leading to underdiagnosis and limiting our understanding of the full neurodevelopmental impact of the disorder (4).

In addition to cognitive and behavioural deficits, FASD patients frequently experience issues with sleep and biological rhythms, as commonly reported by them or their caregivers (5). Simultaneously, sleep disturbances have been reported in other contexts of alcohol exposure, including acute administration, chronic abuse, and dependence, and notably, these disturbances often persist during periods of abstinence (6,7). These observations suggest a link between alcohol exposure, either during perinatal or during adulthood and circadian rhythm and sleep disturbances that have yet to be fully characterized.

Circadian rhythms are endogenous ∼24 h cycles in biological processes that affect various physical, chemical, and behavioural changes in an organism, allowing the adaptation of physiological activities to environmental conditions and homeostasis. In mammals, circadian rhythms are regulated via the master pacemaker located in the suprachiasmatic nucleus (SCN) of the anterior hypothalamus (8). Circadian rhythms can persist in conditions devoid of time cues and are modulated by environmental cycles and internal activity to ensure environmental adaptation (9). This adaptation process is called “entrainment” (10). The main external cue (zeitgeber) that entrains the mammalian circadian system is light (11). Therefore, information captured by the intrinsically photoreceptive retinal ganglion cells (ipRGCs) are transmitted to the SCN through the retino-hypotalamic tract (RHT) (12). Photic information is then transmitted from the SCN to synchronize molecular clocks in central and peripheral tissues.

The molecular clock is cell autonomous and arises from an autoregulatory negative feedback transcriptional network (13). At its core are the main transcriptional activators, CLOCK and its paralogue NPAS2, along with BMAL1, encoded by the *Arntl* gene. CLOCK and NPAS2 can bind to BMAL1 and 2 proteins to form a transcription factor heterodimer. This dimer can activate the transcription of period (*Per*) and cryptochrome (*Cry*) gene families. The resulting PER1-3 and CRY1&2 proteins accumulate and interact to repress the activity of the CLOCK/NPAS2-BMAL1 complex, thereby inhibiting their own transcription in a feedback loop that takes around 24 hours (11,14). Additionally, proteins including nuclear receptors RORα and REV-ERB (encoded by the *Nr1d2* gene) also activate and suppress *Arntl* transcription, respectively, conferring additional stabilization to this feedback system (15,16). Lastly, CLOCK-BMAL1 drives the transcription of D-Box Binding PAR BZIP Factor (DBP), which contributes to the overall regulation of circadian rhythms (14). This delicate and intricate interaction is crucial for the maintenance of the circadian rhythms and homeostasis, as circadian misalignment can compromise physiological systems linked to various neuropathological processes, such as some neurodegenerative diseases (17,18), and other neuropsychiatric conditions such as major depressive disorder (MDD) (19,20) and schizophrenia (21,22).

Alcohol disturbs the organism through various mechanisms affecting cellular and molecular processes, including those governing circadian rhythms (23,24). For instance, experimental animal studies demonstrate that alcohol induces an imbalance between neuronal overactivation and inhibition, impairing dopaminergic, GABAergic, and glutamatergic neurotransmission in key brain regions (25–28). These changes in neurotransmission contribute to behavioural, memory, and cognitive deficits (29–31). Additionally, perinatal and adult alcohol exposure has been shown to alter core body temperature and cortisol rhythmic cycles in rats, causing phase advancements and delays, respectively (32,33). Notably, perinatal alcohol exposure causes enduring alterations in the endogenous rhythmicity of the SCN circadian clock and disrupts clock gene expression in rats, with effects persisting throughout life (34). Such findings suggest that circadian disruption following perinatal alcohol exposure, could potentially underly broader homeostatic dysfunctions and contribute to cognitive and functional impairments observed in prenatal and lactation alcohol exposure (PLAE) rodents. However, the extent of circadian dysregulations caused by perinatal alcohol exposure and their impact on FASD patients remain understudied, and further research is needed to determine how these disturbances may influence deficits across multiple domains in individuals with FASD (35). Moreover, circadian rhythm disruption could serve as a long-term biological marker for PLAE, enhancing diagnostic accuracy for FASD (35).

This study introduces a novel approach to investigate the effects of alcohol exposure on circadian rhythms using the “drinking in the dark” (DID) model as a PLAE procedure in mice, a model previously established by our laboratory (36–38). We sought to investigate whether animals exposed to PLAE displayed alterations of circadian rhythms reflected in locomotor activity alterations during adolescence. In addition, we sought to validate the PLAE as an efficient FASD model by reproducing previously reported spatial memory impairments in this model. Lastly, at a molecular level, we investigated the detrimental effects of PLAE in circadian gene expression across brain areas, including genes from the molecular clock, and other involved in plasticity and cognition, including those related to the expanded endocannabinoid and other neurotransmitter systems.

Overall, our results indicate that PLAE disrupts circadian rhythms in free running conditions during adolescence, impairs spatial memory in adulthood and alters the oscillatory patterns of clock gene expression in a brain region and age dependent manner. Furthermore, we provide evidence that PLAE disrupts circadian expression of genes involved in endocannabinoid signalling, as well as glutamatergic, GABAergic and dopaminergic transmission in areas such as the Hippocampus (HPC) and the Medial Prefrontal Cortex (mPFC). These two areas display a key role in cognitive processes, such as motivation learning, memory, emotional regulation, and motor control (31,39–41), and could be an underlying factor contributing to the pathological FASD phenotype and could potentially serve as early biomarkers of perinatal alcohol exposure.

## METHODS

### Animals

Seven-week-old male (n = 10) and female (n = 20) C57BL/6 breeders were purchased from Charles River (Lyon, France) and transported to the animal facilities (UBIOMEX- PRBB). Breeding began and animals were individually housed in standard cages in a temperature (21° ± 1 °C), humidity (55% ± 10%), and a 12-hour reverse light-dark cycle (lights off from 07:30 to 19:30). The mice were given at least one week to acclimatize to the new conditions before experimentation began during the dark phase under dim red light. Pregnant females were monitored daily for parturition. Date of birth was designated as PD0. After weaning at PD21, the offspring were housed in groups of 4. For experiments with offspring, cohorts were constituted with male and females. Unless specified, food and water were available *ad libitum*. All animal care and experimental procedures adhered to the guidelines set by the European Communities Directive 88/609/EEC for animal research. These procedures received approval from the local ethical committee (CEEA-OH-PRBB) and every effort was made to minimize animal suffering, discomfort, and the number of animals used.

### Drugs

Ethyl alcohol was purchased from Merck Chemicals (Darmstadt, Germany) and diluted in tap water to obtain a 20% (v/v) alcohol solution.

### Drinking in the dark test

The DID procedure in pregnant dams was conducted following the standard protocol previously established in our laboratory (36–38,42–44) to induce PLAE under a binge- like drinking pattern in C57BL/6 mice resulting in blood alcohol concentrations greater than 0.8 g/L (45). For that, 2 days after mating, 20 pregnant females were randomly assigned to either alcohol (n = 10) or water (n = 10) group. The DID was performed during the entire pregnancy and lactation periods for 4 days every week (Figure 1a).

**Figure 1.**
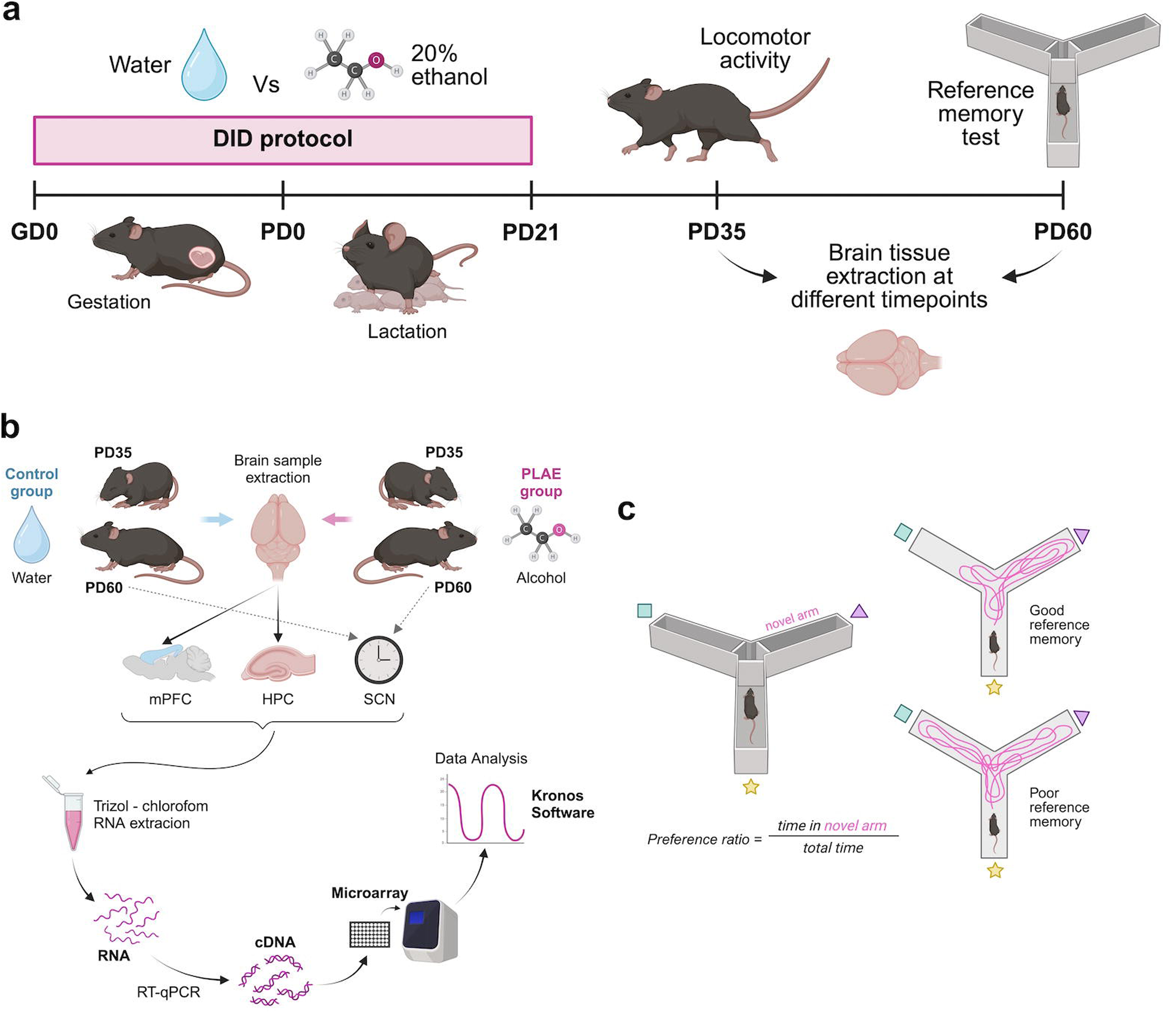
Experimental design. **a** | Schematic illustration of the experimental procedure. **b |** Schematic illustration of experimental design and sample processing. **c |** Schematic representation of the Location Reference Memory test (Y maze). All illustrations are created with Biorender.com. DID, drinking in the dark. GD, gestation day. HPC, hippocampus. mPFC, medial prefrontal cortex. PD, post-natal day. SCN, suprachiasmatic nucleus.

During the first three days, dams had a 2 h access to an alcoholic solution (20% ethanol) or tap water for the Alcohol and Water group respectively. On the fourth day, the access period was extended to 4 h to promote binge drinking. Liquid volumes were recorded before and after every drinking session and expressed as mean ± SEM of the water and alcohol consumption (volume in mL). This protocol continued until pups were weaned at PD21 for approximately 6 weeks, encompassing prenatal and lactation periods as presented in Figure 2a.

**Figure 2.**
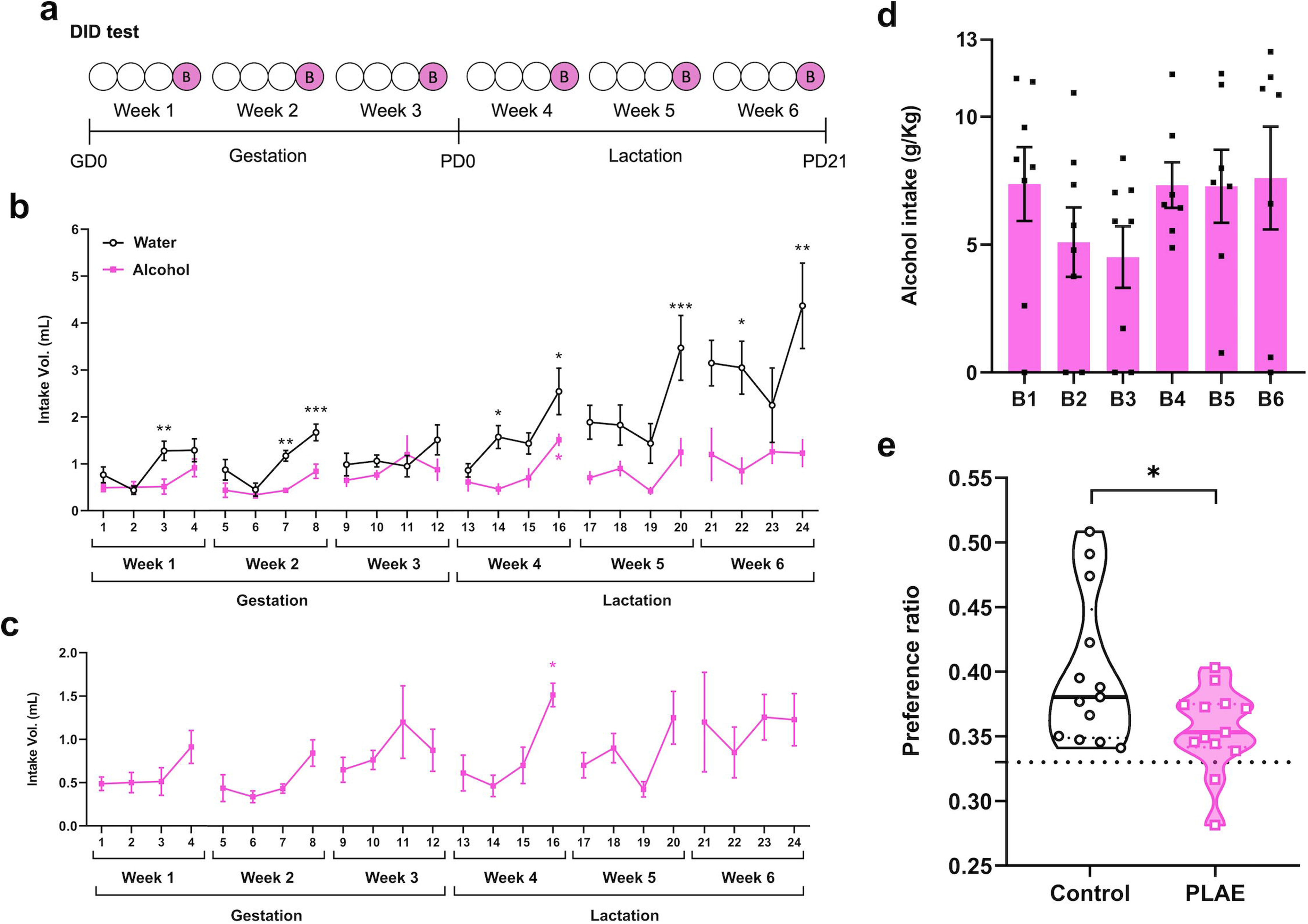
Alcohol consumption during DID test and location memory testing. **a** | Schematic representation of the DID test (Binge 4h). **b |** Volume (mL) of water (n = 8) or alcohol (20% ethanol, n = 8) consumed by the dams during the DID test. Each point represents the mean ± SEM. Two-way ANOVA with repeated measures revealed significant differences between water and alcohol assigned groups. Black asterisks show alcohol vs. water differences (same day), pink show differences across alcohol days, *p < .05, ** p < .01, ***p < .001. **c |** Volume (mL) of alcohol consumed by the dams during the DID test (n = 8). Each point represents the mean ± SEM. Significant differences revealed by a one-way ANOVA with repeated measures are indicated when appropriate, *p < 0.05. **d |** Alcohol intake (g of alcohol / kg of body weight) in the six sessions of binge-like drinking. Each bar represents the mean ± SEM (n = 8). **e |** Performance of PLAE (alcohol, n =13) vs Control (water, n=13) PD60 mice in the Location Reference Memory test (Y-maze) was assigned using a two-way ANOVA, indicating a statistically significant difference with *p < .05 between experimental groups. Data are expressed in violin plots as mean, individual values and 25 and 75% percentiles. Dotted line at y = 0.33 represents chance-level performance. DID, drinking in the dark. GD, gestation day. PD, post-natal day. PLAE, prenatal and lactation alcohol exposure.

### Experimental design

Starting from the first day of gestation, pregnant dams were subjected to the DID protocol, explained above. After weaning at PD21, pups were separated into two experimental groups: water-exposed (Control) and alcohol-exposed (PLAE) mice (Figure 1a). First, we assessed behavioural parameters related to circadian and cognitive deficits. For this, at PD35, a cohort of animals (n_Control_ = 5, n_PLAE_ = 5) was subjected to a study of locomotor activity under free-running conditions. At PD60, a second cohort of animals was used to perform the location reference memory test.

For the gene expression assessment, two batches of animals from both experimental groups were sacrificed to obtain tissue samples from the mPFC and HPC (PD35 and PD60) and from the SCN (PD60). Sample collection was performed at six different time points to evaluate changes in gene expression throughout the light-dark circle. Samples were then processed as described in section “Tissue collection and RNA extraction” (Figure 1b).

### Locomotor activity

Locomotor activity was evaluated in offspring to assess circadian rhythms under free running conditions. Starting at PD35, animals (n = 10, 5 females and 5 males) were individualized and their home cages placed inside a frame featuring infra-red technology that allowed for the measuring of activity counts in intervals of 30 minutes (LE8825, LE8816; Panlab s.l.u., Barcelona, Spain). Baseline activity was analysed during a two-day acclimatization period under a L/D cycle (12 hours of continuous light followed by 12 hours of continuous darkness) with zeitgeber time 0 (ZT0) indicating the onset of the light phase (8:00 AM). The results from the two days of basal conditions were averaged for more effective analysis, corresponding to the condition L/D category. Then, locomotion was recorded over a period of 6 days under constant darkness (D/D). Environmental conditions, including light intensity during the light phase and ambient temperature, were maintained uniformly throughout the experiment. See Figure 1a for a schematic representation of the experimental design.

To objectively compare the daily rhythms of the two experimental groups, the data were processed to determine the oscillatory nature of the locomotor activity and to reconstruct sinusoidal curves for each day’s activity for each experimental group. This allowed precise evaluation of the differences in daily rhythmicity between groups throughout the experiment.

### Location reference memory test

The location reference memory test was performed in offspring, at PD60 to evaluate spatial memory (n = 13 for both groups) as previously described (38). A Y-maze with three identical arms separated by 120° angles each featuring one visual cue was employed (Figure 1c). The experiment included one training and one test session with a one-hour inter-trial interval. During training, one of the arms was blocked and labelled as the novel arm. Animals were introduced in a randomly assigned arm and allowed to freely explore the two available corridors for 5 minutes. During the test, the blockage was removed, and mice were allowed to freely explore all arms for 5 minutes. The Smart Software (Panlab S.L.U., Barcelona, Spain) was used to track the location and the movement of the animals across the maze. The preference ratio was calculated using the following equation:

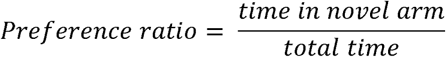

### Tissue collection and RNA extraction

Control and PLAE mice were euthanized by cervical dislocation at PD35 and PD60 at six different time points ZT2, 6, 10, 14, 18 and 22, as previously described (46). The offspring were distributed equally across the different sacrifice points to avoid biases derived from maternal alcohol consumption. Their brains were quickly removed and the HPC, mPFC, and the SCN were dissected using a 1 mm brain matrix. Samples were immediately stored at -80 °C until later processing.

Total RNA was extracted and isolated from brain tissue using TRIzol^TM^ reagent (Invitrogen, 15596026) and isopropanol as previously described (47). RNA concentration was determined with Thermo Scientific™ NanoDrop™ One/OneC. RNA aliquots were stored at -80°C until reverse transcription (RT) (Applied Biosystems™ 4374967) was performed to synthesize cDNA following the manufacturer’s instructions. Total RNA from SNC samples was adjusted to 100 ng/uL before the RT, while total RNA from mPFC and HIP samples were adjusted to 200 ng/uL. cDNA aliquots of all samples were stored at -20 °C awaiting further processing (Figure 1b).

### OpenArray^TM^ technology

To perform the gene expression analyses, custom OpenArray^TM^ plates were designed and purchased from Thermo Fisher Scientific (Table S1). For this, 2.5 μL of cDNA sample was combined with 2.5 μL TaqMan OpenArray^TM^ Real-Time Master Mix (Thermo Fisher #4462159) and loaded into a single well of a 384-well plate (48). Custom OpenArray^TM^ plates were then automatically loaded using the AccuFill System (AccuFill System User Guide, PN4456986) and run in Applied Biosystems™ QuantStudio™ 12K Flex Real-Time PCR. Amplification of the sequence of interest was normalized to reference endogenous genes, specifically, the geometric mean of actin f3 (*Actb*), f32 microglobulin (*B2m*), hypoxanthine-guanine phosphoribosyl transferase (*Hprt1*), and glyceraldehyde-3-phosphate dehydrogenase (*Gapdh*). Data were analysed with the ThermoFisher ExpressionSuite Software and fold-change values were calculated using the ΔΔCT method (49) (Figure 1b) with Control ZT02 values as reference sample.

### Statistical analysis

Data was first thoroughly analysed for normality and heteroscedasticity. When these assumptions were met, parametric tests were used to evaluate statistical differences. On this basis, data obtained from the DID test were analysed using two-way analysis of variance (ANOVA) with group (Control and PLAE) as a between-subject factor and day as a within-subject factor, followed by Bonferroni post hoc comparisons. One-way ANOVAs with repeated measures were used to analyse alcohol intake during the DID test and locomotor activity, followed by Dunnet’s post hoc testing. Data from the location reference memory test were analysed using a two-way ANOVA. All data are represented as the mean ± SEM and significance was set at *p* < 0.05. These analyses and the corresponding graphs were performed using GraphPad Prism version 8.0 (GraphPad Software. San Diego, California, USA).

Biological rhythms of locomotor activity and target genes was determined using the R (2024.04.2+764) (50) package Kronos, (https://github.com/thomazbastiaanssen/kronos). Gene expression data were normalized to Control group values at ZT02 to account for baseline differences across experimental conditions. Statistical analyses were computed using the base R stats package. Outliers from gene expression datasets were removed if deviated more than 1.5 × IQR below the 1^st^ quartile or above the 3^rd^ quartile (51). For all tests a significance threshold of a = 0.05 was used.

Additionally, Biorender.com was employed to create some of the illustrations, all appropriately cited in the figure footnotes. All code used to analyse data from this study is freely available at the GitHub repository https://github.com/mariareinacampos/Circadian-Rhythms-and-PLAE under the GNU license.

## RESULTS

### Maternal alcohol consumption during gestation and lactation

Liquid volumes consumed during the DID protocol were measured before and after each drinking session and expressed as mean ± SEM of water and alcohol consumption (in mL) over the 6-week period (Figure 2b). A two-way ANOVA revealed significant differences in intake volumes between water and alcohol throughout the protocol (Figure 2b, Table S2). There was a significant increase in water consumption during weeks 4, 5 and 6, corresponding to the lactation process, in comparison to alcohol consumption.

A one-way ANOVA with repeated measures was used to evaluate alcohol intake differences within each drinking session (Figure 2c, Table S3). In week 4, there was a significant effect of day on alcohol consumption (F(3,31) = 7.897, *p* = 0.0012). Dunnett’s post hoc comparisons indicated significant increases in alcohol intake on binge day 4 compared to days 1 (p = 0.0170), 2 (p = 0.0059) and 3 (p = 0.0238) of the same DID session. No significant differences were observed during the other weeks.

Additionally, the grams of alcohol consumed per kilogram of mouse body weight were calculated for each binge session (Figure 2d). One-way ANOVA with repeated measures found no significant differences in the amount of alcohol consumed between the binge sessions throughout the protocol (F(5,47) = 0.9139, p = 0.4460), assuming blood levels of intoxication on binge days as previously described (38).

### Spatial memory impairments in the PLAE group

A location reference memory test was conducted at PD60 to evaluate the spatial working and reference memory. Results showed that the mean preference ratio of both groups was significantly higher than chance level (0.33). A two-way ANOVA revealed no significant effects of sex (F (1, 22) = 0.1259, *p* = 0.7261) or the interaction between sex and the experimental groups (Control vs. PLAE) (F (1, 22) = 0.2881, *p* = 0.5968).

However, there was a significant main effect of group (F(1, 22) = 5.378, p = 0.0301), indicating that mice exposed to alcohol during the prenatal and lactation periods (PLAE group, n = 13) showed impaired spatial memory performance in the reference test compared to mice exposed to water (Control group, n = 13) in adulthood (PD60) (Figure 2e). Since the validating experiment did not reveal significant differences between sexes, all subsequent analyses were performed on pooled data, without considering sex as an independent factor.

### PLAE mice show altered locomotor activity

Spontaneous locomotor activity of mice (n = 5 per group) was monitored during adolescence (PD35) in dark-dark conditions (D/D) for a period of 6 consecutive days to assess daily rhythms. Representative actograms of both experimental groups are shown in Figure 3a and 3b. When analysed for circadian rhythmicity, the constructed sinusoidal curves (Table S4) reveal an apparent amplitude increment between the days of the experiment for both groups, although a one-way ANOVA for repeated measures analysis revealed that this increase was not significant for either Control (F (6, 34) = 1.882, *p* = 0.2064) or PLAE (F (6, 34) = 2.68, *p* = 0.1223) mice. To further investigate the potential increase in activity, total daily movement counts were analysed throughout the experiment (Figure 3e and 3f). A one-way repeated-measures ANOVA revealed a significant progressive increase in activity exclusively in the PLAE group (F (6, 34) = 10.66, *p* = 0.0055) with post hoc testing reporting significant differences between the baseline condition and the final day of the experiment on the 6^th^ day (*p* = 0.0140). This increase in number of activity counts from baseline to last day of free running was not present in the Control group.

**Figure 3.**
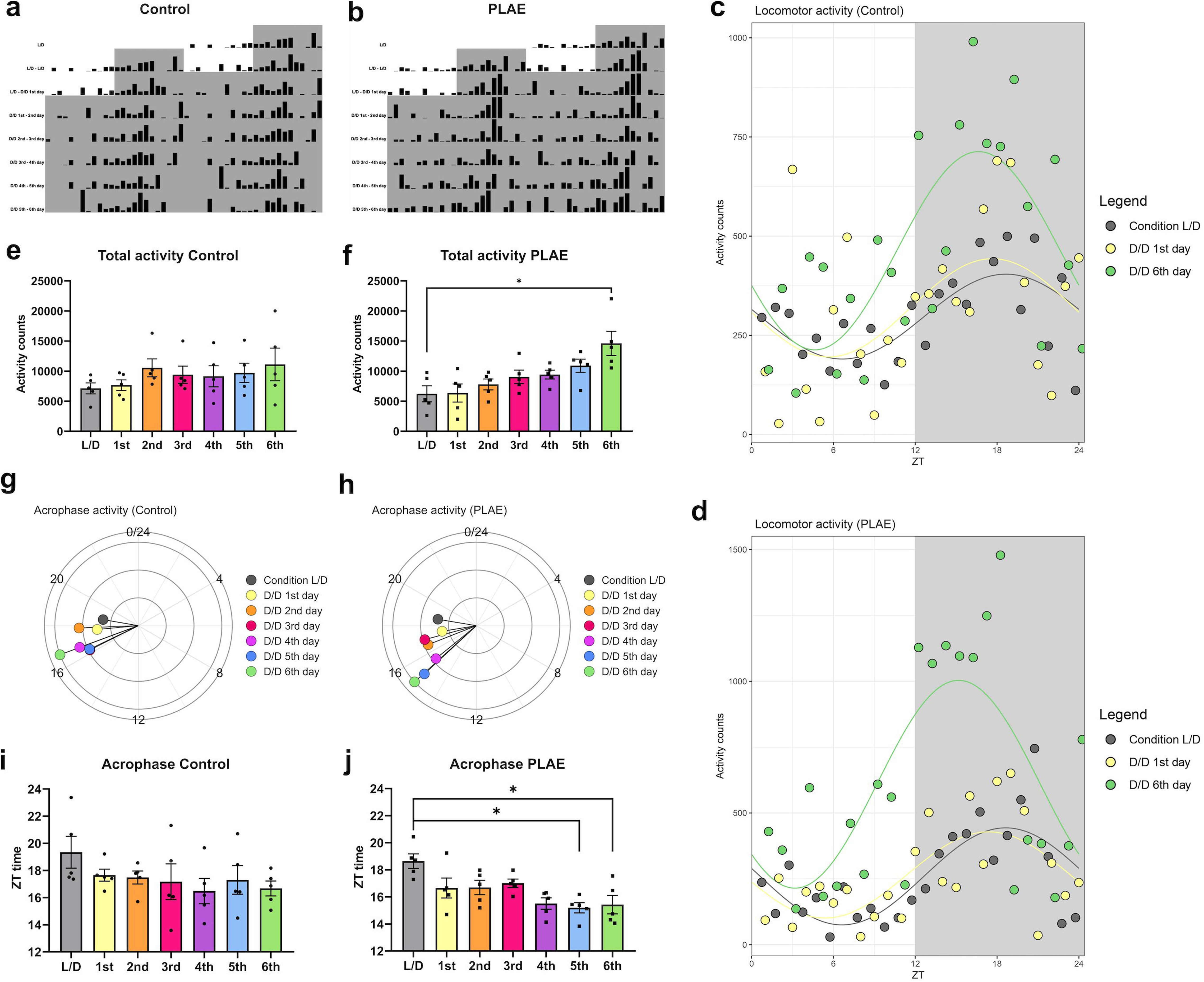
Free running locomotor activity patterns in Control and PLAE mice. **a, b** | Representative actograms for Control (a) and PLAE (b) mice, double plotted over 48 hours. Each bar indicates the number of movements made per hour. **c, d |** Sinusoidal plots of daily locomotor activity for Control (c) (n = 5) and PLAE (d) (n = 5) mice of basal conditions (Condition L/D), D/D 1^st^ day and D/D 6^th^ day of the experiment. L/D condition represents the mean activity during two days of light-dark cycles. n = 5. **e, f |** Total daily activity counts throughout the experiment for Control (e) and PLAE (f). Significant differences are found in the PLAE group the basal L/D condition and day 6 under D/D conditions (*p < 0.05, One-way ANOVA with repeated measures). Each bar represents the average amplitude ± SEM, with n = 5. **g, h |** Circular diagrams showing the daily acrophase of activity for Control (g) and PLAE (h) with n = 5. **i, j |** Bar plot of activity acrophase time for each experimental day for Control (i) and PLAE (j) groups. In PLAE mice, significant differences were found between the basal L/D condition and days 5 and 6 under D/D conditions (*p < 0.05, One-way repeated measures ANOVA). Each bar represents the average acrophase ± SEM, with n = 5. D/D, dark-dark. L/D, light-dark. PLAE, prenatal and lactation alcohol exposure. ZT, zeitgeber time.

In addition, the acrophase values of locomotor activity were analysed for each day of the experiment. As shown in Figures 3g and 3h, both experimental groups exhibited an apparent phase advancement of the peak of maximum activity over the course of the study. Although acrophase values were identical between groups under baseline conditions, the PLAE group exhibited a significant decrease in acrophase by the end of the experiment compared to its initial value. A one-way repeated-measures ANOVA confirmed a progressive reduction in acrophase time for the PLAE group (F (6, 34) = 8.706, p = 0.0033), with post hoc tests revealing significant differences between the basal condition (L/D) and days 5 (p = 0.0261) and 6 (p = 0.0188) (Figure 3j). In contrast, the same analysis in the Control group did not yield statistically significant differences (Figure 3i).

### PLAE mice exhibit loss of circadian rhythmicity in both circadian and non- circadian clock genes in mPFC and HPC

Alcohol exposure during neurodevelopmental stages appears to exert a consistent impact upon the gene expression across brain regions (mPFC and HPC) and age (PD35 and PD60), as reflected by similar percentages of genes with altered rhythmicity (loss or modification) (Figure 4a). Further details are provided in Figure 4b, which lists all genes exhibiting an oscillatory expression pattern under any of the experimental conditions. This analysis highlighted that the genes affected by alcohol include not only those involved in the molecular regulation of circadian rhythms but also those associated with other physiological functions, such as expanded endocannabinoid system, several neurotransmitters, plasticity, and motivation. Notably, these alterations are observed across all two evaluated brain regions and at both ages studied.

**Figure 4.**
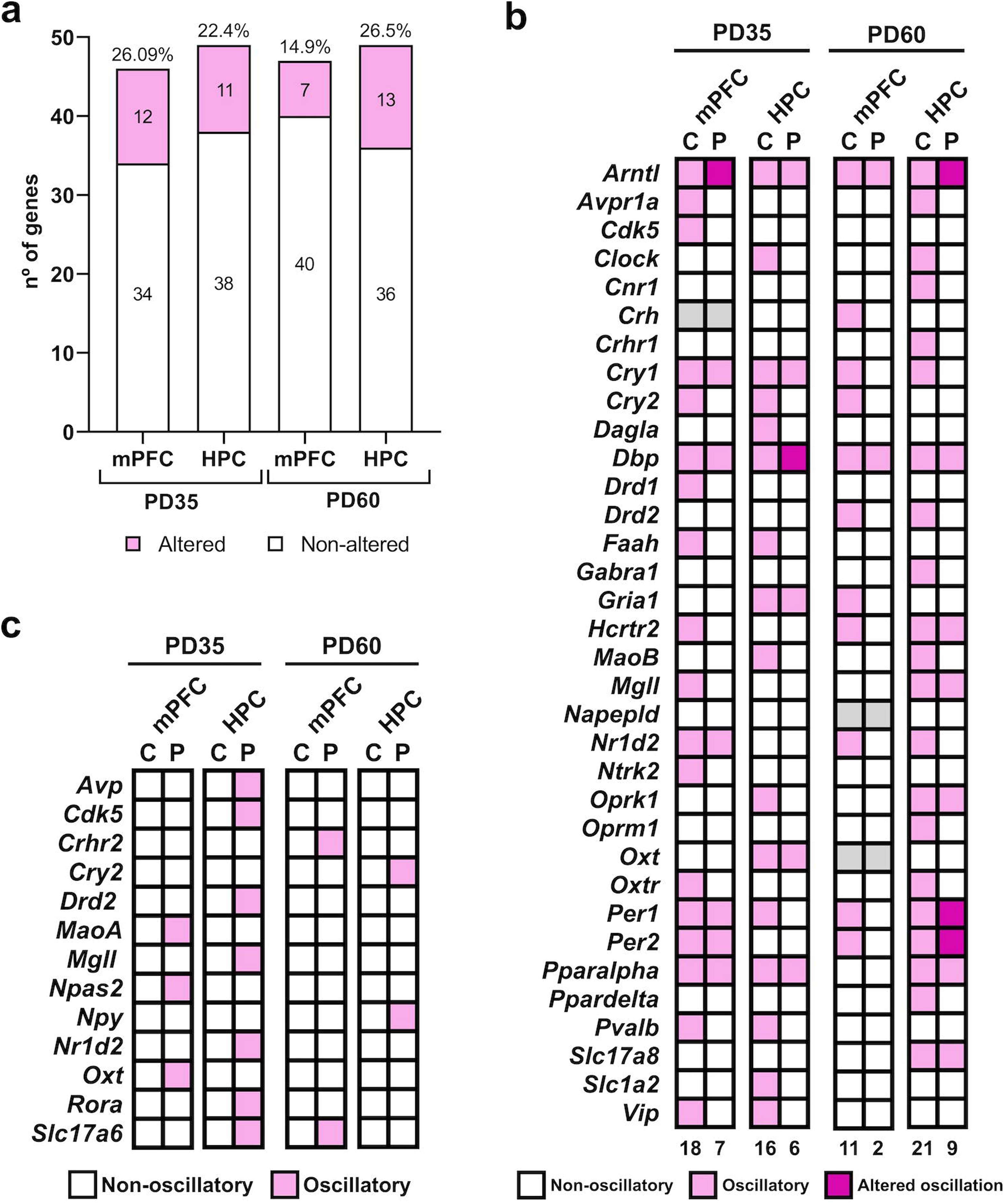
Oscillatory characterization of gene array. **a** | Number of modulated genes in mPFC and HPC at PD35 and PD60. Bars represent the total number of genes analysed in each region, with the percentage of altered genes (loss or change in rhythmicity) indicated above each bar and the number of genes of each category indicated inside the corresponding bar. **b |** Heatmap (C = Control, P = PLAE). Genes that were not oscillatory in any condition are not shown. Grey cells indicate missing data for these genes at certain brain regions. Numbers below indicate the number of oscillatory genes per each column. **c |** Heatmap (C = Control, P = PLAE) indicating de novo cycling genes detected in each brain region of interest. HPC, Hippocampus. mPFC, Medial Prefrontal Cortex. PD, post-natal day.

The mPFC showed 26.09% of altered genes (loss or change in rhythmicity) during adolescence (PD35) and 14.9% in adulthood (PD60), with distinct sets of affected genes at each age (Figure 4a). At PD35, the altered clock genes are *Arntl* and *Cry2* (Table S5), while non-clock genes include *Avpr1a*, *Cdk5*, *Drd1*, *Faah*, *Hcrtr2*, *MgII*, *Ntrk2*, *Oxtr*, *Pvalb*, and *Vip* (Table S6). At PD60, the altered clock genes shift to *Cry1*, *Cry2*, *Nr1d2*, and *Per1/2* (Table S5), while non-clock genes include *Drd2*, *Gria1*, and *Hcrtr2* (Table S7).

The HPC also demonstrated age-dependent changes in gene expression, with 22.4% of altered genes at PD35 and 26.5% at PD60 (Figure 4a). At PD35, the affected clock genes include *Clock*, *Cry2*, and *Per1* (Table S5), and non-clock genes such as *Dagla*, *Faah*, *MaoB*, *Oprk1*, *Pvalb*, *Slc1a2*, and *Vip* (Table S8). By PD60, the altered clock genes include *Arntl*, *Clock*, *Cry1*, and *Nr1d2* (Table S5), with non-clock genes such as *Avpr1a*, *Cnr1*, *Crhr1*, *Drd2*, *Gabra1*, *MaoB*, *Oprm1*, *Oxtr*, and *Ppard* (Table S9). Moreover, all regions exhibited a small subset of genes that, while lacking an oscillatory pattern in Control conditions, display *de novo* cycling under PLAE conditions (Figure 4c, Table S10).

Lastly, focusing on the circadian rhythm-related genes included in the array, a subset was selected due to their relevance in the control of biological clock, and their expression patterns are presented in Figure 5a. These graphs reveal that *Arntl*, the key gene in circadian regulation, experiences a phase delay of up to 5 hours under PLAE conditions compared to basal conditions. This delay in peak *Arntl* expression is more pronounced at PD35 (Figure 5b and 5c). Moreover, this acrophase shift appears to be specific to *Arntl*, as other circadian clock genes, such as *Dbp* and *Per1* (Figure 5a, Table S5) show either identical acrophase values across conditions or lose their rhythmicity entirely under PLAE.

**Figure 5.**
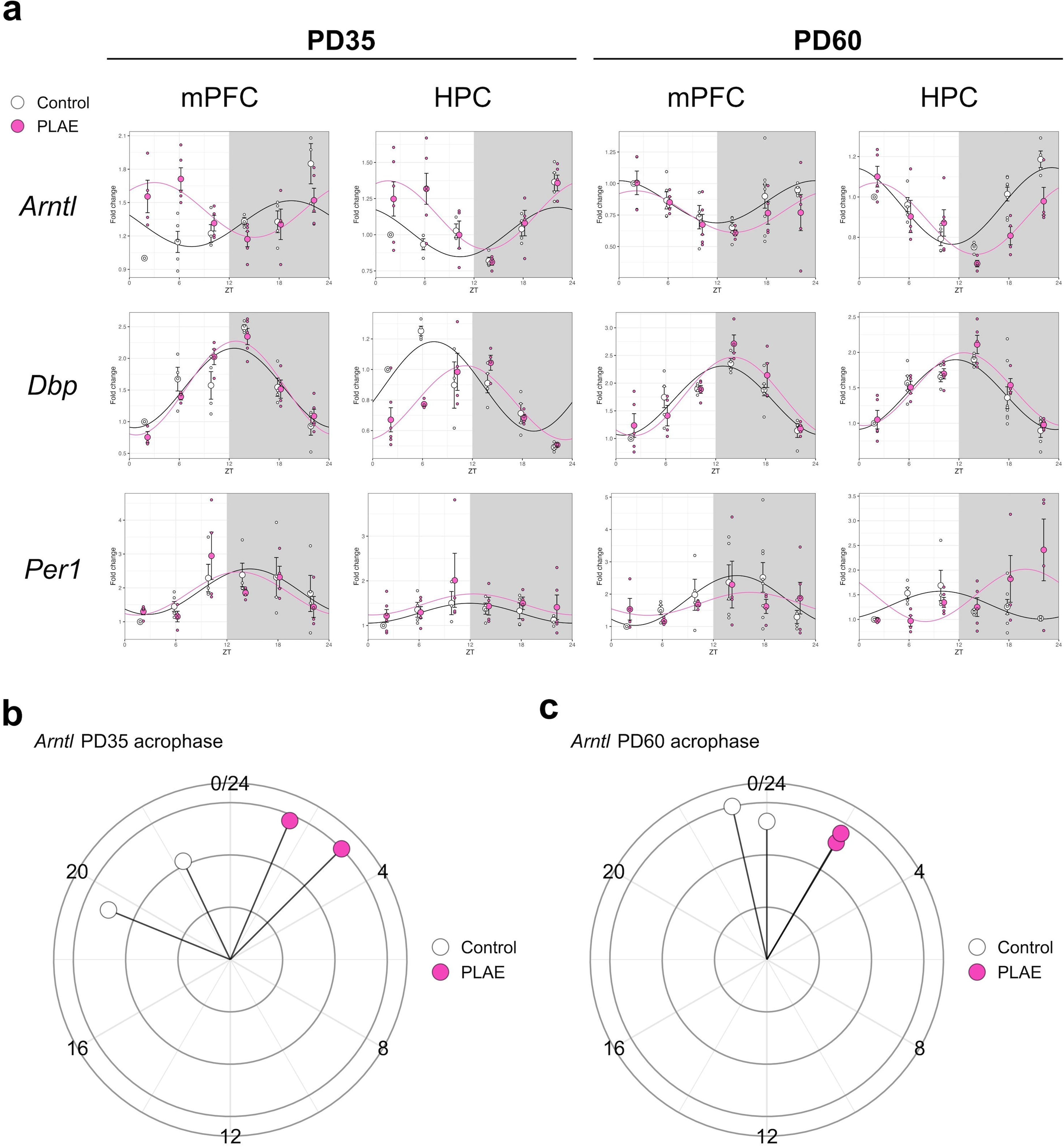
Oscillatory transcription of clock genes. **a** |Sinusoidal plots of the relative gene expression of *Arntl, Dbp*, and *Per1* showing relative gene expression (normalized to Control ZT02) upon water (Control) and alcohol (PLAE) intake, each value represents the mean ± SEM with n = 3-5 for each timepoint/treatment. **b, c |** Circular diagrams showing Arntl acrophase for each combination of group, brain area and mice age. HPC, Hippocampus. mPFC, Medial Prefrontal Cortex. PD, post-natal day. ZT, zeitgeber time.

### Clock gene expression in the SCN

As illustrated in Figure 6a, most clock genes lose their oscillatory expression pattern, with the notable exceptions of *Bmal1* and *Dbp*. A closer examination of the expression patterns for *Arntl*, *Dbp*, and *Per1* in Figure 6b reveals that the delay in *Arntl* acrophase, previously observed in the specific clocks (mPFC and HPC) at both PD35 and PD60, is absent in the SCN. This finding is further supported by data shown in Figure 6c (Table S11), which demonstrates the lack of the phase shift previously detected in specific clocks.

**Figure 6.**
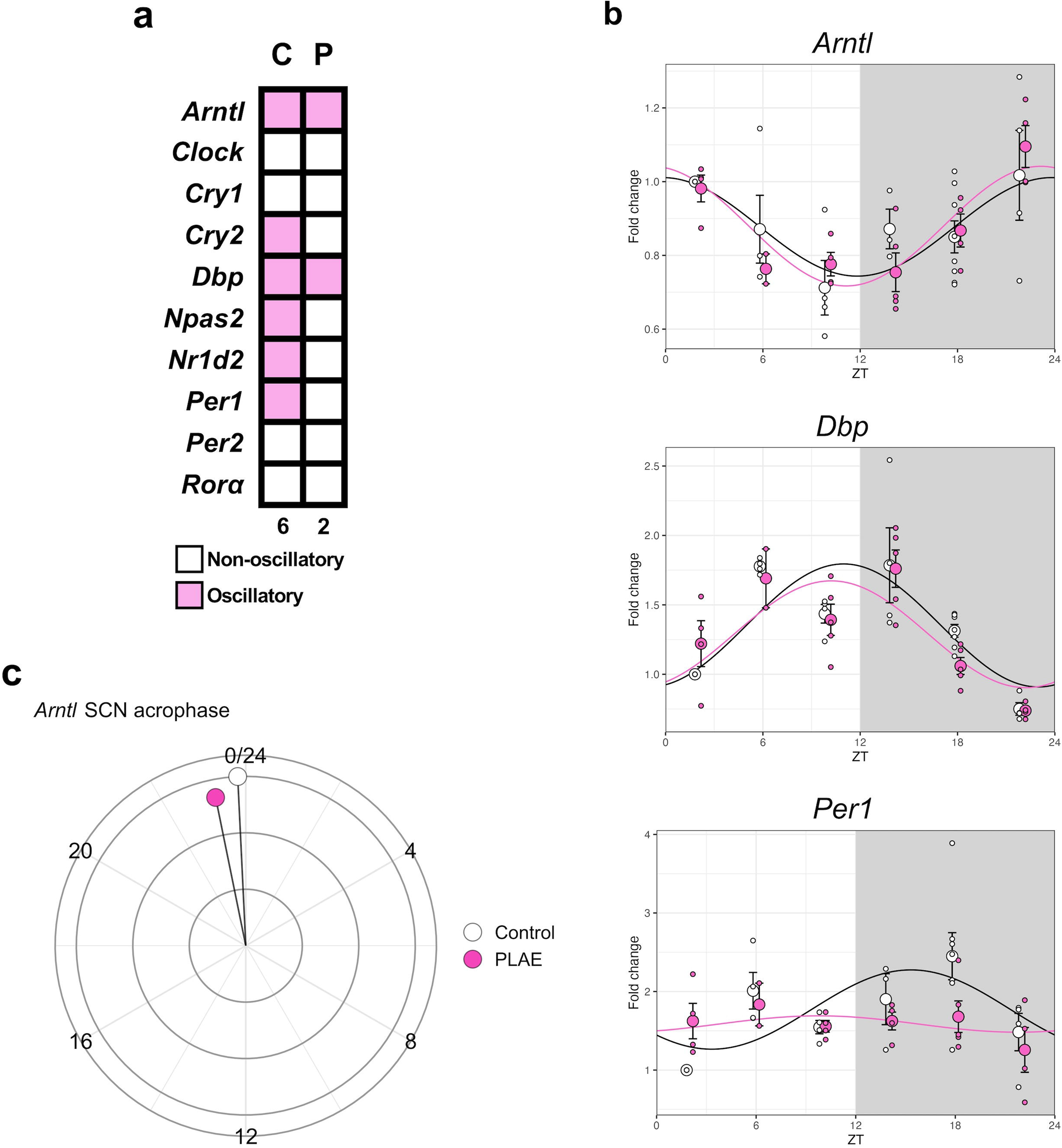
Clock gene expression in the SCN at PD60. **a** |Heatmap (C = Control, P = PLAE) showing oscillatory or non-oscillatory gene expression of main clock genes in the SCN at PD60. **b |** Sinusoidal plots of the relative gene expression of *Bmal1, Dbp*, and *Per1* showing relative gene expression (normalized to Control ZT02) upon water (Control) and alcohol (PLAE) intake, each value represents the mean ± SEM with n = 3-5 for each timepoint/treatment. **c |** Circular diagram showing Arntl (Bmal1) acrophase in the SCN at PD60 of Control and PLAE groups. PD, post-natal day. SCN, Suprachiasmatic Nucleus.

## DISCUSSION

The results of this study provide evidence that PLAE induces persistent effects on the genetic regulation of clock genes in the offspring, disrupting the oscillatory behaviour of both clock-related and non-clock-related genes. Additionally, these genetic alterations are accompanied by impairments in spontaneous locomotor activity and adult location reference memory, suggesting that changes in the biological rhythms might contribute to the behavioural and cognitive disruptions observed in our PLAE animal model and that are also present in FASD patients. Our study covers the period of adolescence and early adulthood, showing alterations in gene expression in both periods, although we cannot rule out that the alterations extend to other periods of life.

Alcohol consumption during the DID protocol followed a pattern consistent with previous results (36,44,52). Water consumption was significantly higher than alcohol intake during the final weeks of the protocol, corresponding to the lactation period. This difference was likely driven by an increased fluid demand associated with milk production (53). Our results indicate that the PLAE group exhibited significantly impaired location reference memory compared to the Control group. This result is in accordance with previous observations in the same experimental conditions in our laboratory (37,38).

An important manifestation of altered circadian rhythms in rodents is the disruption in locomotor activity, also present in FASD (2). Adolescence is one of the key periods of life in which FASD symptomatology is identified, as we have previously reported (38). The analysis of the oscillatory pattern of activity curves reveals that PLAE mice exhibit a progressive phase advancement, starting their activity period earlier each day of the experiment. These findings in mice are consistent with a previous study in which rats exposed to alcohol during the early postnatal period (PD4-PD9) exhibited a significantly shorter circadian period in wheel-running activity compared to controls under D/D conditions (54). To the best of our knowledge, this is the first time these alterations have been demonstrated in mice. Therefore, our findings contribute to the growing body of literature suggesting that alcohol exposure during the prenatal and neonatal period can lead to long-term alterations in circadian behaviour, particularly a reduction in the circadian period of activity rhythms. Additionally, our study shows that PLAE mice exhibit higher locomotor activity counts after six days in free running conditions in comparison to their basal locomotor activity under standard light cycles. This finding is consistent with the results from a previous study in which rats were exposed to alcohol during the end of the gestational period (34). Observing this consistent phenotype across animal models is particularly relevant because FASD patients are often misdiagnosed as Attention Deficit and Hyperactivity Disorder (ADHD), being the hyperactive behaviour a hallmark symptom (55–57). Therefore, the elevated activity observed in PLAE mice further validates this model as a reliable representation of behaviours commonly associated with FASD.

Although it has been already proven that oscillatory gene expression, such as *Per1/2*, emerges in the SCN as early as embryonic day 18 in mice (58), circadian rhythms continue to develop and refine throughout the lifespan of an organism (59). Our findings suggest that the establishment of circadian rhythms is a developmental process, in which adolescence plays a key role in its organization (60). However, in our study, we did not observe an increase in the number of genes that have an oscillatory pattern of expression between adolescent (PD35) and young adult (PD60) brain samples. The lack of increased oscillatory gene expression may indicate that the rhythmicity of our genes of interest is already well-established by adolescence. Although we observed some genes that exhibited a sinusoidal expression pattern at PD35, some failed to maintain their oscillatory patterns in the same brain regions at PD60. This raises important questions about the stability of oscillatory patterns at PD35. It remains unclear whether these oscillations at this stage are transient, representing an intermediate developmental phase, or if they reflect stable circadian rhythms that are disrupted in PLAE. Given the lack of studies examining the developmental trajectory of oscillatory gene expression beyond the SCN, further research is needed to determine whether these oscillations would naturally stabilize or shift with age and whether PLAE alters this development.

Our findings highlight a novel aspect of PLAE-related neurodevelopmental alterations, emphasizing that its effects extend beyond the SCN to genes involved in cognitive, emotional, and motivational processes, the main domains altered in FASD (2,61). The fact that these genes specifically lose their oscillatory expression under PLAE further supports the idea that this disruption is not merely a transient developmental phase but rather a direct consequence of perinatal alcohol exposure. This reinforces the hypothesis that PLAE interferes with the stability of gene rhythmicity in regions critical for neurobehavioral function, potentially contributing to the persistent cognitive and emotional deficits observed in FASD. Interestingly, in our study, we also observed the emergence of *de novo* cycling genes under alcohol exposure. Although this is a relatively novel area of research, *de novo* cycling has been previously reported in a circadian rhythm study (62). This study, which examined the role of circadian rhythms in response to chronic stress, found increased *de novo* cycling in the HPC of stress- resilient mice, the same region where we identified most of the *de novo* cycling genes in our study. While further research is needed to determine whether these newly emerging rhythms are beneficial or contribute to the long-term deficits observed in FASD, this parallel suggests that *de novo* cycling may function as a compensatory mechanism in response to PLAE-induced disruptions, potentially serving as an adaptive response to mitigate its neurodevelopmental impact.

To better understand how these molecular disruptions contribute to the FASD phenotype, it is essential to analyse the physiological functions of the genes that lose their oscillatory expression following PLAE. One of the affected systems is the endocannabinoid system (ECS), which plays a critical role in synaptic plasticity and learning process but also emotional and motivational behaviours (40) that have been reported to be altered in PLAE (38). Our findings reveal that key components involved in endocannabinoid synthesis (*Dagla* and *Napepld*) and degradation (*MgII* and *Faah*), alongside the canonical CB1 receptor (*Cnr1*) (63), exhibit a disrupted rhythmicity under PLAE conditions. The fact that crucial players in the ECS exhibit expression impairments by premature alcohol exposure is consistent with previous results reported in our lab (38) and findings from other authors that report that perinatal alcohol exposure affects brain development via an ECS-related mechanism that contributes to the long-lasting learning and memory deficits in rodents (64). Beyond the ECS, PLAE also disrupts the rhythmic expression of opioid receptors (*Oprm1*, *Oprk1*), which are involved in motivation, reward processing, and emotional regulation (65). These receptors interact closely with the ECS (66) and are implicated in behavioural domains significantly affected in FASD (37,42).

Additionally, the altered expression of corticotropin-releasing hormone (*Crh*) and its receptor (*Crhr1*) could underlie the heightened anxiety susceptibility reported in FASD patients (67). The oxytocin receptor (*Oxtr*) also exhibits altered rhythmicity, potentially contributing to the social deficits, characteristic of FASD (61,68,69).

Furthermore, PLAE-induced disruptions extend to genes regulating major neurotransmitter systems, including glutamate (*Gria1*, *Slc17a8*, *Slc1a2*), dopamine (*Drd1*, *Drd2*, *Ppa1r1b*) and GABA (*Gabra1*). These neurotransmitters are essential for learning, memory, emotional regulation, and motor control, playing a crucial role in maintaining homeostatic responses and adaptive behaviour (70). The dysregulation of these systems may further explain the persistent neurobehavioral impairments observed in FASD.

Interestingly, a study on mouse embryonic fibroblast cells demonstrated that the deletion of *Cdk5* lengthened the circadian period, suggesting that this kinase plays a role in circadian synchronization *in vitro* but contradicting findings from *in vivo* mouse models (71). In our study, we observed that *Cdk5* exhibited oscillatory expression in the mPFC at PD35 but lost this rhythmicity in the PLAE group. Given that *Cdk5* is altered precisely at this developmental stage, our findings also contrast with the *in vitro* results, as PLAE mice exhibited a significant acrophase advancement, indicating a shortened circadian period.

One of the most significant findings from the molecular analysis was the apparent loss of activity integration between the master pacemaker, the SCN, and specific clocks in regions such as the mPFC and HPC. This is supported by the sinusoidal expression of *Arntl* under PLAE, which exhibits a delay of several hours in the mPFC and HPC but remains constant and unaltered in the SCN. A previous study similarly reported alcohol-induce phase advancements of *Per1*, another key clock gene, in peripheral clocks, while the SCN remained unaffected (33), reinforcing the hypothesis that prenatal alcohol might play a role in disrupting connectivity and functional communication between brain regions (43).

Moreover, the observed *Arntl* dysregulation aligns with findings showing that knocking out *Arntl* in cortical neurons does not affect SCN rhythms but significantly disrupts circadian clock machinery in the neocortex and HPC (72). This suggests that the circadian disturbances observed in PLAE result from both alcohol-induced disruptions in interregional synchronization and the direct consequences of altered *Arntl* expression on brain function. These findings further support the hypothesis that the SCN loses its ability to properly synchronize specific brain clocks under PLAE, likely due to alcohol-induced synaptic disruptions (43), while *Arntl* dysregulation itself contributes to cognitive and emotional deficits, exacerbating PLAE-related impairments.

In conclusion, this study presents a novel approach to investigating circadian rhythm impairments in a validated PLAE mouse model, revealing subtle yet widespread alterations caused by alcohol exposure during early brain development. The gaps in knowledge about these molecular and neuronal processes underscore the critical need for continued research into circadian rhythms under both baseline conditions and perturbations, such as prenatal alcohol exposure. Our findings demonstrate significant effects on spatial memory, alterations in free running locomotor activity patterns, and disruptions in the oscillatory behaviour of both clock and non-clock genes, including genes involved in the expanded endocannabinoid system (e.g. *Cnr1, Dagla, Faah*) and other neurotransmitter systems (e.g. *Oprm1, Slc17a8, Drd1, Gabra1*). By advancing the understanding of the intricate relationship between early alcohol exposure and biological circadian rhythms, this study paves the way for developing targeted therapies to alleviate the neurobehavioral challenges faced by FASD patients and to facilitate earlier and more accurate diagnoses.

## ACKNOWLEDGEMENTS

This work was supported by the Ministerio de Ciencia e Innovación, “Grant PID2022-136962OB-100 - MCIN/AEI/10.13039/501100011033 and by ERDF A way of making Europe”, Ministerio de Sanidad (Delegación del Gobierno para el Plan Nacional sobre Drogas #2023/005 and #Exp2022/008695 Fondos de Recuperación, Transformación y Resiliencia (PRTR) Union Europea, and by the Generalitat de Catalunya, AGAUR (#2021SGR00485). OV is recipient of an ICREA Academia Award (Institució Catalana de Recerca i Estudis Avançats, Generalitat de Catalunya).

## AUTHOR CONTRIBUTIONS

MR-C, IG-L and OV conceptualized and designed the study. IG-L, and MR-C conducted the behavioural experiments. Data analysis was performed by MR-C, IG-L and MM. The manuscript was written by MR-C, MM and OV. IG-L, MM and OV provided critical revisions. OV supervised the project. All authors reviewed and approved the final version of the manuscript.

## SUPPLEMENTARY INFORMATION

- Table S1. Assay IDs and the corresponding targets of the OpenArray^TM^ Thermo Fisher.
- Table S2. Two-way ANOVA with repeated measures results for water or alcohol intake during the DID protocol.
- Table S3. One-way ANOVA with repeated measures results for alcohol intake during the DID protocol.
- Table S4. Kronos’ rhythmic analysis output of locomotor activity in mice exposed to water (Control) or alcohol (PLAE).
- Table S5. Kronos rhythmic analysis output of clock genes in medial prefrontal cortex and hippocampus of mice exposed to water (Control) or alcohol (PLAE).
- Table S6. Kronos rhythmic analysis output of genes in medial prefrontal cortex at PD35 of mice exposed to prenatal water (Control) or alcohol (PLAE).
- Table S7. Kronos rhythmic analysis output of genes in medial prefrontal cortex at PD60 of mice exposed to water (Control) or alcohol (PLAE).
- Table S8. Kronos rhythmic analysis output of genes in hippocampus at PD35 of mice exposed to water (Control) or alcohol (PLAE).
- Table S9. Kronos rhythmic analysis output of genes in hippocampus at PD60 of mice exposed to water (Control) or alcohol (PLAE).
- Table S10. Kronos rhythmic analysis output of de novo cycling genes in medial prefrontal cortex and hippocampus of mice exposed to water (Control) or alcohol (PLAE).
- Table S11. Kronos rhythmic analysis output of clock genes in suprachiasmatic nucleus at PD60 of mice exposed to water (Control) or alcohol (PLAE).

## ABBREVIATURES

*Actb*: Actin f3
ADHD: Attention deficit and hyperactivity disorder
ANOVA: Analyses of variance
*B2m*: f32 microglobulin
CB1: Endocannabinoid receptor *Cnr1*
Crh: Corticotropin-releasing hormone
*Crhr1*: Corticotropin-releasing hormone receptor
D/D: Dark-dark
DBP: D-Box PAR BZIP factor
DID: Drinking in the Dark
ECS: Endocannabinoid system
FAS: Foetal alcohol syndrome
FASD: Foetal alcohol spectrum disorders
*Gapdh*: Glyceraldehyde-3-phosphate dehydrogenase
HPC: Hippocampus
*Hprt1*: Hypoxanthine-guanine phosphoribosyl transferase
ipRGCs: Intrinsically photoreceptive retinal ganglion cells
L/D: Light-dark
MDD: Major depressive disorders
mPFC: Medial Prefrontal Cortex
RHT: Retino-Hypotalamic tract
*Oxtr*: Oxytocin receptor
PD: Post-natal day
PLAE: Prenatal and lactation alcohol exposure
SCN: Suprachiasmatic nucleus
ZT: Zeitgeber time

## REFERENCES

1. Chudley AE, Conry J, Cook JL, Loock C, Rosales T, LeBlanc N. Fetal alcohol spectrum disorder: Canadian guidelines for diagnosis. CMAJc: Canadian Medical Association Journal, 2005, 172: S21.

2. Mattson SN, Crocker N, Nguyen TT. Fetal Alcohol Spectrum Disorders: Neuropsychological and Behavioral Features. Neuropsychol Rev, 2011 21(2):101.

3. Lange S, Probst C, Gmel G, Rehm J, Burd L, Popova S. Global Prevalence of Fetal Alcohol Spectrum Disorder Among Children and Youth: A Systematic Review and Meta-analysis. JAMA Pediatr, 2017, 171(10):956.

4. Nash K, Rovet J, Greenbaum R, Fantus E, Nulman I, Koren G. Identifying the behavioural phenotype in Fetal Alcohol Spectrum Disorder: sensitivity, specificity and screening potential. Arch Womens Ment Health, 2006, 9(4):181–6.

5. Wengel T, Hanlon-Dearman AC, Fjeldsted B. Sleep and sensory characteristics in young children with fetal alcohol spectrum disorder. J Dev Behav Pediatr, 2011, 32(5): 384–92.

6. Colrain IM, Turlington S, Baker FC. Impact of Alcoholism on Sleep Architecture and EEG Power Spectra in Men and Women. Sleep, 2009, 32(10):1341–52.

7. Brower KJ. Insomnia, alcoholism and relapse. Sleep Med Rev., 2003, 7(6):523–39.

8. Tamura EK, Oliveira-Silva KS, Ferreira-Moraes FA, Marinho EAV, Guerrero-Vargas NN. Circadian rhythms and substance use disorders: A bidirectional relationship. Pharmacol Biochem Behav., 2021, 201:173105.

9. Refinetti R. Integration of biological clocks and rhythms. Compr Physiol, 2012, 2(2):1213–39.

10. Jiménez A, Lu Y, Jambhekar A, Lahav G. Principles, mechanisms and functions of entrainment in biological oscillators. Vol. 12, Interface Focus. Royal Society Publishing; 2022. p. 20210088.

11. Mohawk JA, Green CB, Takahashi JS. Central and peripheral circadian clocks in mammals. Annu Rev Neurosci., 2012, 35: 445–62.

12. Do MTH, Yau KW. Intrinsically Photosensitive Retinal Ganglion Cells. Physiol Rev., 2010, 90(4):1581.

13. Lowrey PL, Takahashi JS. Mammalial circadian biology: Elucidating genome-wide levels of temporal organization. Annu Rev Genomics Hum Genet, 2004, 5: 441.

14. Takahashi JS. Transcriptional architecture of the mammalian circadian clock. Nat Rev Genet., 2016, 18(3): 164–79.

15. Sato TK, Panda S, Miraglia LJ, Reyes TM, Rudic RD, McNamara P, et al. A functional genomics strategy reveals rora as a component of the mammalian circadian clock. Neuron, 2004, 43(4):527–37.

16. Preitner N, Damiola F, Luis-Lopez-Molina, Zakany J, Duboule D, Albrecht U, et al. The orphan nuclear receptor REV-ERBα controls circadian transcription within the positive limb of the mammalian circadian oscillator. Cell, 2002, 110(2):251–60.

17. Wulff K, Gatti S, Wettstein JG, Foster RG. Sleep and circadian rhythm disruption in psychiatric and neurodegenerative disease. Nat Rev Neurosci., 2010, 11(8):589–99.

18. Canever JB, Queiroz LY, Soares ES, de Avelar NCP, Cimarosti HI. Circadian rhythm alterations affecting the pathology of neurodegenerative diseases. J Neurochem, 2023, 168(8):1475–89.

19. Manosso LM, Duarte LA, Martinello NS, Mathia GB, Réus GZ. Circadian Rhythms and Sleep Disorders Associated to Major Depressive Disorder: Pathophysiology and Therapeutic Opportunities. CNS & Neurological Disorders-drug Targets, 2023, 23(9): 1085–100.

20. Amidfar M, Garcez ML, Kim YK. The shared molecular mechanisms underlying aging of the brain, major depressive disorder, and Alzheimer’s disease: The role of circadian rhythm disturbances. Prog Neuropsychopharmacol Biol Psychiatry 2023, 123: 110721.

21. Waite F, Sheaves B, Isham L, Reeve S, Freeman D. Sleep and schizophrenia: From epiphenomenon to treatable causal target. Schizophr Res. 2020, 221:44–56.

22. Bromundt V, Köster M, Georgiev-Kill A, Opwis K, Wirz-Justice A, Stoppe G, et al. Sleep-wake cycles and cognitive functioning in schizophrenia. Br J Psychiatry, 2011, 198(4): 269–76.

23. Gulick D, Gamsby JJ. Racing the clock: The role of circadian rhythmicity in addiction across the lifespan. Pharmacol Ther., 2018, 188:124–39.

24. Meyrel M, Rolland B, Geoffroy PA. Alterations in circadian rhythms following alcohol use: A systematic review. Prog Neuropsychopharmacol Biol Psychiatry, 2020, 99: 109831–109831.

25. Avchalumov Y, Mandyam CD. Synaptic Plasticity and its Modulation by Alcohol. Brain Plasticity [Internet]. 2020, 6(1): 111.

26. Hughes BA, Bohnsack JP, O’Buckley TK, Herman MA, Morrow AL. Chronic Ethanol Exposure and Withdrawal Impairs Synaptic GABAA Receptor-Mediated Neurotransmission in Deep Layer Prefrontal Cortex. Alcohol Clin Exp Res., 2019, 43(5): 832.

27. Tateno T, Robinson HPC. The mechanism of ethanol action on midbrain dopaminergic neuron firing: a dynamic-clamp study of the role of I(h) and GABAergic synaptic integration. J Neurophysiol., 2011, 106(4): 1901–22.

28. Lovinger DM, White G, Weight FF. NMDA receptor-mediated synaptic excitation selectively inhibited by ethanol in hippocampal slice from adult rat. J Neurosci., 1990, 10(4):1379.

29. McNab F, Varrone A, Farde L, Jucaite A, Bystritsky P, Forssberg H, et al. Changes in cortical dopamine D1 receptor binding associated with cognitive training. Science, 2009, 323(5915): 800–2.

30. Sarter M, Bruno JP, Parikh V. Abnormal Neurotransmitter Release Underlying Behavioral and Cognitive Disorders: Toward Concepts of Dynamic and Function-Specific Dysregulation. Neuropsychopharmacology, 2007, 32(7): 1452–61.

31. McEntee WJ, Crook TH. Glutamate: its role in learning, memory, and the aging brain. Psychopharmacology (Berl), 1993, 111(4):391–401. Available from: https://link.springer.com/article/10.1007/BF02253527

32. Handa RJ, Zuloaga DG, McGivern RF. Prenatal ethanol exposure alters core body temperature and corticosterone rhythms in adult male rats. Alcohol. 2007, 41(8): 567–75.

33. Guo R, Simasko SM, Jansen HT. Chronic Alcohol Consumption in Rats Leads to Desynchrony in Diurnal Rhythms and Molecular Clocks. Alcohol Clin Exp Res., 2016, 40(2): 291–300.

34. Sakata-Haga H, Dominguez HD, Sei H, Fukui Y, Riley EP, Thomas JD. Alterations in circadian rhythm phase shifting ability in rats following ethanol exposure during the third trimester brain growth spurt. Alcohol Clin Exp Res., 2006, 30(5): 899–907.

35. Inkelis SM, Thomas JD. Sleep in Infants and Children with Prenatal Alcohol Exposure. Alcohol Clin Exp Res., 2018, 42(8): 1390–405.

36. Cantacorps L, Alfonso-Loeches S, Moscoso-Castro M, Cuitavi J, Gracia-Rubio I, López-Arnau R, et al. Maternal alcohol binge drinking induces persistent neuroinflammation associated with myelin damage and behavioural dysfunctions in offspring mice. Neuropharmacology. 2017 Sep 1;123:368–84.

37. Cantacorps L, Montagud-Romero S, Luján MÁ, Valverde O. Prenatal and postnatal alcohol exposure increases vulnerability to cocaine addiction in adult mice. Br J Pharmacol . 2020, 177(5):1090–105.

38. Garcia-Baos A, Pastor A, Gallego-Landin I, de la Torre R, Sanz F, Valverde O. The role of PPAR-γ in memory deficits induced by prenatal and lactation alcohol exposure in mice. Mol Psychiatry, 2023, 28(8): 3373–83.

39. Campolongo P, Trezza V. The endocannabinoid system: a key modulator of emotions and cognition. Front Behav Neurosci. 2012, 6:73.

40. Zanettini C, Panlilio L V., Alicki M, Goldberg SR, Haller J, Yasar S. Effects of Endocannabinoid System Modulation on Cognitive and Emotional Behavior. Front Behav Neurosci., 2011, 5: 57.

41. Gepshtein S, Li X, Snider J, Plank M, Lee D, Poizner H. Dopamine Function and the Efficiency of Human Movement. J Cogn Neurosci., 2013, 26(3): 657.

42. Esteve-Arenys A, Gracia-Rubio I, Cantacorps L, Pozo OJ, Marcos J, Rodríguez-Árias M, et al. Binge ethanol drinking during adolescence modifies cocaine responses in mice. J Psychopharmacol., 2017, 31(1): 86– 95.

43. Cantacorps L, González-Pardo H, Arias JL, Valverde O, Conejo NM. Altered brain functional connectivity and behaviour in a mouse model of maternal alcohol binge-drinking. Prog Neuropsychopharmacol Biol Psychiatry. 2018, 84: 237–49.

44. García-Baos A, Puig-Reyne X, García-Algar Ó, Valverde O. Cannabidiol attenuates cognitive deficits and neuroinflammation induced by early alcohol exposure in a mice model. Biomedicine & Pharmacotherapy. 2021, 141: 111813.

45. National Institute on Alcohol Abuse and Alcoholism (NIAAA). Alcohol’s Effects on Health. Drinking Levels Defined, 2023.

46. Berbegal-Sáez P, Gallego-Landin I, Macía J, Alegre-Zurano L, Castro-Zavala A, Welz PS, et al. Lack of Bmal1 leads to changes in rhythmicity and impairs motivation towards natural stimuli. Open Biol., 2024, 14(7): 240051.

47. Castro-Zavala A, Martín-Sánchez A, Montalvo-Martínez L, Camacho-Morales A, Valverde O. Cocaine-seeking behaviour is differentially expressed in male and female mice exposed to maternal separation and is associated with alterations in AMPA receptors subunits in the medial prefrontal cortex. Prog Neuropsychopharmacol Biol Psychiatry, 2021, 109: 110262.

48. Abraham NA, Campbell AC, Hirst WD, Nezich CL. Optimization of small- scale sample preparation for high-throughput OpenArray analysis. J Biol Methods, 2021, 8(1): e143.

49. Livak KJ, Schmittgen TD. Analysis of Relative Gene Expression Data Using Real-Time Quantitative PCR and the 2−ΔΔCT Method. Methods. 2001, 25(4): 402–8.

50. R Core Team. R: A Language and Environment for Statistical Computing. Vienna, Austria: R Foundation for Statistical Computing; 2023.

51. Kwak SK, Kim JH. Statistical data preparation: management of missing values and outliers. Korean J Anesthesiol, 2017, 70(4): 411.

52. Rhodes JS, Best K, Belknap JK, Finn DA, Crabbe JC. Evaluation of a simple model of ethanol drinking to intoxication in C57BL/6J mice. Physiol Behav. 2005, 84(1): 53–63.

53. Hydration as a limiting factor in lactation. American Journal of Biological Anthropology, 1998, 10(2): 151–61.

54. Allen GC, West JR, Chen WJA, Earnest DJ. Neonatal Alcohol Exposure Permanently Disrupts the Circadian Properties and Photic Entrainment of the Activity Rhythm in Adult Rats. Alcohol Clin Exp Res., 2005, 29(10): 1852.

55. Raldiris TL, Bowers TG, Towsey C. Comparisons of Intelligence and Behavior in Children With Fetal Alcohol Spectrum Disorder and ADHD. J Atten Disord. 2018; 22(10):959–970.

56. Peadon E, Elliott EJ. Distinguishing between attention-deficit hyperactivity and fetal alcohol spectrum disorders in children: clinical guidelines. Neuropsychiatr Dis Treat., 2010, 6: 509–15.

57. Burger PH, Goecke TW, Fasching PA, Moll G, Heinrich H, Beckmann MW, et al. How does maternal alcohol consumption during pregnancy affect the development of attention deficit/hyperactivity syndrome in the child. Fortschr Neurol Psychiatr., 2011, 79(9): 500–6.

58. Landgral D, Koch CE, Oster H. Embryonic development of circadian clocks in the mammalian suprachiasmatic nuclei. Front Neuroanat., 2014, 8(DEC):119895.

59. Olejniczak I, Pilorz V, Oster H. Circle(s) of Life: The Circadian Clock from Birth to Death. Biology, 2023, 12(3): 383.

60. Carmona-Alcocer V, Abel JH, Sun TC, Petzold LR, Doyle FJ, Simms CL, et al. Ontogeny of Circadian Rhythms and Synchrony in the Suprachiasmatic Nucleus. J Neurosci., 2018, 38(6): 1334.

61. Kodituwakku PW. Neurocognitive Profile In Children With Fetal Alcohol Spectrum Disorders. Dev Disabil Res Rev., 2009, 15(3): 224.

62. Savva C, Vlassakev I, Bunney BG, Bunney WE, Massier L, Seldin M, et al. Resilience to Chronic Stress Is Characterized by Circadian Brain-Liver Coordination. Biological Psychiatry Global Open Science, 2024, 4(6): 100385.

63. Cristino L, Bisogno T, Di Marzo V. Cannabinoids and the expanded endocannabinoid system in neurological disorders. Nature Reviews Neurology 2019, 16(1): 9–29.

64. Basavarajappa BS. Fetal Alcohol Spectrum Disorder: Potential Role of Endocannabinoids Signaling. Brain Sci., 2015, 5(4): 493.

65. van Steenbergen H, Eikemo M, Leknes S. The role of the opioid system in decision making and cognitive control: A review. Cogn Affect Behav Neurosci 2019, 19(3):435–58.

66. Ahmad T, Lauzon NM, de Jaeger X, Laviolette SR. Cannabinoid Transmission in the Prelimbic Cortex Bidirectionally Controls Opiate Reward and Aversion Signaling through Dissociable Kappa Versus μ-Opiate Receptor Dependent Mechanisms. J. Neurosci., 2013, 33(39): 15651.

67. Wilhoit LF, Scott DA, Simecka BA. Fetal Alcohol Spectrum Disorders: Characteristics, Complications, and Treatment. Community Ment Health J., 2017, 53(6): 711–8.

68. Borroto-Escuela DO, Cuesta-Marti C, Lopez-Salas A, Chruścicka-Smaga B, Crespo-Ramírez M, Tesoro-Cruz E, et al. The oxytocin receptor represents a key hub in the GPCR heteroreceptor network: potential relevance for brain and behavior. Front Mol Neurosci., 2022, 15: 1055344.

69. Pierzynowska K, Gaffke L, Żabińska M, Cyske Z, Rintz E, Wiśniewska K, et al. Roles of the Oxytocin Receptor (OXTR) in Human Diseases. Int J Mol Sci., 2023, 24(4): 3887.

70. Takei N, Momiyama T, Ueda S, Kaibuchi K, Tsuboi D, Nagai T, et al. Neuromodulator regulation and emotions: insights from the crosstalk of cell signaling. Front Mol Neurosci., 2024, 17:1376762.

71. Ripperger JA, Chavan R, Albrecht U, Brenna A. Physical Interaction between Cyclin-Dependent Kinase 5 (CDK5) and Clock Factors Affects the Circadian Rhythmicity in Peripheral Oscillators. Clocks Sleep, 2022, 4(1):201.

72. Bering T, Carstensen MB, Wörtwein G, Weikop P, Rath MF. The Circadian Oscillator of the Cerebral Cortex: Molecular, Biochemical and Behavioral Effects of Deleting the Arntl Clock Gene in Cortical Neurons. Cereb Cortex, 2018, 28(2):644–57.

